# A social bee (*Bombus terrestris*) learns to associate a novel odor with social context

**DOI:** 10.1101/2025.05.30.657092

**Authors:** Etya Amsalem, Abraham Hefetz

## Abstract

Many pheromone responses are hardwired into the insect’s nervous system and are essential for critical behaviors such as mating, alarm signaling, and trail following. Workers of social species are assumed to respond innately to queen pheromones, affecting their behavior and reproductive physiology. However, accumulating evidence suggests that learning and experience may also shape responses to social chemical cues. To examine if the response to queen-associated odors can be learned, we exposed *Bombus terrestris* workers to a novel odor by treating the queen daily with unfamiliar floral scents, either anisyl alcohol or methyl anthranilate. We allowed perfumed queens to establish colonies and examined worker attraction and egg laying following exposure to these odors in the absence of the queen either with or without brood as context. In a two-choice olfactometer, workers preferred the queen odor they learned during colony development. Exposure of workers to the queen-associated odor in the absence of the queen influenced worker egg laying only in the presence of brood, surprisingly resulting in increased egg laying. Our study demonstrates that workers can learn novel odors that are associated with their queen or nest, which in turn function as context-dependent social cues that modify their social and reproductive behavior.

## Introduction

Social insect life is largely regulated by pheromones, which are especially important for maintaining reproductive division of labor between fecund queens and sterile workers. Such pheromones are thought to regulate worker fertility and promote the formation of cohesive social groups where most females refrain from reproduction and act as helpers. Despite extensive research, only a handful of reproduction-regulating pheromones have been identified in social insects ^1–3^. This limitation is due to several factors, such as the overemphasis on the queen as the main driver of reproductive inhibition in the colony ^1,4^ while neglecting contextual elements ^5^, and the assumption that innate pheromonal responses are inflexible. This narrow framing can limit the scope of discovery and underrepresent the diversity of pheromonal systems, leading researchers to overlook the complexity and flexibility of pheromonal communication. Indeed, many pheromonal responses are innate: male moths instinctively respond to pheromones released by conspecific females without prior exposure ^6^, ants follow pheromone trails instinctively to locate food or return to the nest ^7^, and honeybees respond to isopentyl acetate alarm pheromone component that elicits a stinging defense behavior in face of a threat ^8^.

However, a growing body of evidence from both vertebrates and invertebrates shows that responses to pheromones can be shaped by learning, and modified or overridden by experience ^9^. When reared in a foster nest, stingless bees of the species *Scaptotrigona subobscuripennis* follow the trail pheromones of their foster colony rather than those of their natal colony ^10^, male fruit flies are less attracted to females marked with cis-vaccenyl acetate, a pheromone transferred during mating that signals a female has already mated, and their attraction is further suppressed through learning after rejection by a mated female ^11,12^, honeybee workers modify their response to Nasonov gland pheromone (geraniol and citral) from appetitive to aversive when they learn to associate it with an electric shock ^13^, butterflies are able to learn a new ratio of their sex pheromone blend ^14^, and butterfly larvae not only learn a new host odor after exposure, but can also transfer their preference to their offspring ^15^.

While many insect species are capable of learning ^16,17^, and pheromonal responses can be altered through experience, innate responses to pheromones regulating reproduction and social behavior may be especially subject to modification, given the potential fitness costs of responding inappropriately to these signals. Queen pheromones in eusocial insects, for example, regulate worker reproduction by either manipulating or providing information that workers use to adjust their own reproductive behaviors ^18^, and they differ from other pheromones in several ways that can increase flexibility in pheromonal response. First, queen pheromones are competitive in nature ^19^ and therefore more susceptible to cheating. Such pheromones often serve to resolve reproductive conflicts between the queen, who benefits from being the sole reproducer, and the workers, who in some cases benefit from laying their own eggs ^20^. Given the high fitness consequences for workers, additional forms of assurance may be required to convey reliable information about the queen’s fecundity and functionality, regardless of the mechanism by which queen pheromones operate. Second, pheromones regulating reproduction operate in a highly context-dependent manner ^5^. For example, *Bombus impatiens* workers ignore the queen pheromones unless they are presented alongside young brood and/or visual cues from the queen^21^. Context reinforces signal reliability and provides environmental relevance, and may also enable sensory integration, which may be essential for eliciting a response and enhancing overall effectiveness ^5,22,23^. Because context varies, responses to reproduction-mediating pheromones may be more easily shaped by learning and experience than other pheromonal systems. Indeed, it has been suggested that behaviors specific to time and place are more likely to depend on learning ^16^.

Here, we examined whether the innate response to queen pheromone in *Bombus terrestris* can be modified by learning. Bumblebees are an excellent species to examine the flexibility in pheromonal response since they are annual social bees exhibiting large flexibility in reproductive behavior ^24^. Colonies are established by a single mated queen, develop through the spring and the summer, eventually producing sexuals (i.e., new queens and males) and decline in the fall, during which only the mated queens survive and enter a winter diapause ^24^.

Following colony establishment, the young queen starts producing workers, and fully controls their ovarian activation and egg laying, but approximately one month later, during a “competition phase” ^25^ workers start producing males from unfertilized eggs and engage in aggressive behavior towards each other and the queen. If workers are separated from the queen (queenless, QL), they activate their ovaries and lay eggs as a function of age: within 7-9 days if newly emerged ^26^ and within 2-3 days if older^27^. Brood also affect worker reproduction: in the presence of young larvae, workers still activate their ovaries but lay fewer eggs, whereas in the presence of pupae they lay more eggs ^27–30^.Reproductive division of labor in bumblebee colonies is assumed to be regulated by various means, including context dependent queen pheromones ^21,31^, the young brood ^27,29,30,32^, signals produced by workers ^33–36^ and other sociobiological factors including density ^37^ and the queen’s background ^38^. Resorption of eggs, but not ovary activation, in *B. terrestris* was reported to be influenced by pentacosane (C25) ^39^, a component abundant on both the queen and worker cuticle, as well as in various exocrine glands ^31,33,35,36,40^. However, later studies challenged this finding, showing no gene expression differences in *B. terrestris* workers exposed to C25 or solvent control ^41^, no reproductive effects in response to C25 or related hydrocarbons in *B. impatiens* ^42,43^, and a meta-analysis revealing no difference in cuticular hydrocarbon profile complexity between social and solitary species ^44^. Despite the debate about the specific chemical compound regulating worker reproduction, it has been consistently shown that the queen’s presence and her chemical secretions affect worker reproduction ^21,45,46^.

To test whether exposure to queen-associated novel odors affects worker behavior and reproductive physiology, we treated young founder queens daily by applying an external odor, either anisyl alcohol (AA) or methyl anthranilate (MA), throughout colony development. As colonies matured, workers were tested for behavioral preference between the odor they had experienced during development (hereafter, the ‘familiar odor’) or an alternative odor (hereafter the ‘alternative odor’). We further tested the reproductive output of workers in the absence of the queen following daily exposure to either the familiar or alternative odors, either in the absence or presence of brood from their original colony (as a contextual signal). We hypothesized that if workers could learn to associate the externally applied odor with their queen or nest, they would be preferably attracted to the familiar odor as well as alter their reproductive physiology in its presence. We further hypothesized that the presence of a contextual signal (brood) would enhance the workers’ response to the externally applied odor and increase reproductive inhibition.

## Methods

### Bees

Colonies of *Bombus terrestris* (n=8) were obtained from Poliam, Yad Mordechai, Israel on May 2024, a few days after the emergence of the first worker. All colonies were kept in incubators in darkness, 28° C and 50% RH, and bees were provided with a sucrose solution and fresh pollen purchased from Poliam, Yad Mordechai. At the experiment onset, all colonies contained a queen, 4-7 workers, and a small amount of brood at various stages that were housed in large wooden cages throughout the experiment (Supplementary Information, Fig. S1A). From that point onward, queens were treated daily with 1 mg/1 µl of either Anisyl alcohol (AA) or Methyl anthranilate (MA). Both compounds are liquids at room temperature and were used in their natural, solvent-free form (4 colonies per treatment). The solution was applied to the queen’s thorax using a pipette inserted into the colony with a minimum disturbance to the queens. Colony size was monitored daily, and sampling of workers (see below under “experimental design”) started once the colony reached 20 workers. Thereafter, for 35 days from the onset of the experiment, colony size was maintained at or near 20 workers. Treatment in all colonies started at least one week before the first workers were sampled and continued daily. Colonies were used as long as no signs of competition were observed. i.e., no worker reproduction or aggression, no multiple egg cells open at the same time and no presence of gyne larvae have been observed ^24^.

### Experimental design

The experiments examining the effect of applied odors on worker reproduction were conducted either without or in the presence of brood (experiments 1 and 2, respectively, Fig. 1). We further conducted behavioral assays using a T-maze during the first experiment (see below). In experiment 1 workers were sampled progressively between days 12 to 30, whereas in experiment 2, all workers were sampled on days 31-35. In both experiments, nestmate workers of random age were sampled in groups of 3 and placed in a small plastic cage for 7 days (Supplementary Information, Fig. S1B). Callow workers, indicated by their silvery appearance, were not sampled. Thus, all workers had at least 2-3 days to interact with their scented queen prior to sampling. Colonies and cages were kept in separate incubators according to the odor treatment to prevent the exposure of volatiles of the alternative odor. Cages that were assigned a solvent control treatment were evenly split between the two incubators. All treatments were applied within the same time window, i.e., every morning between 9 to 11 am. In total, 171 cages of 3 workers were set in experiment 1 and 77 in experiment 2. Mortality of a single bee occurred in 14 cages (0.05% of the bees) and was randomly distributed across treatments. All these cages were discarded, resulting in a total of 160 and 74 cages in experiments 1 and 2, respectively. The split of sample size per colony and treatment is provided in Supplementary Information, Table S1.

**Figure 1.**
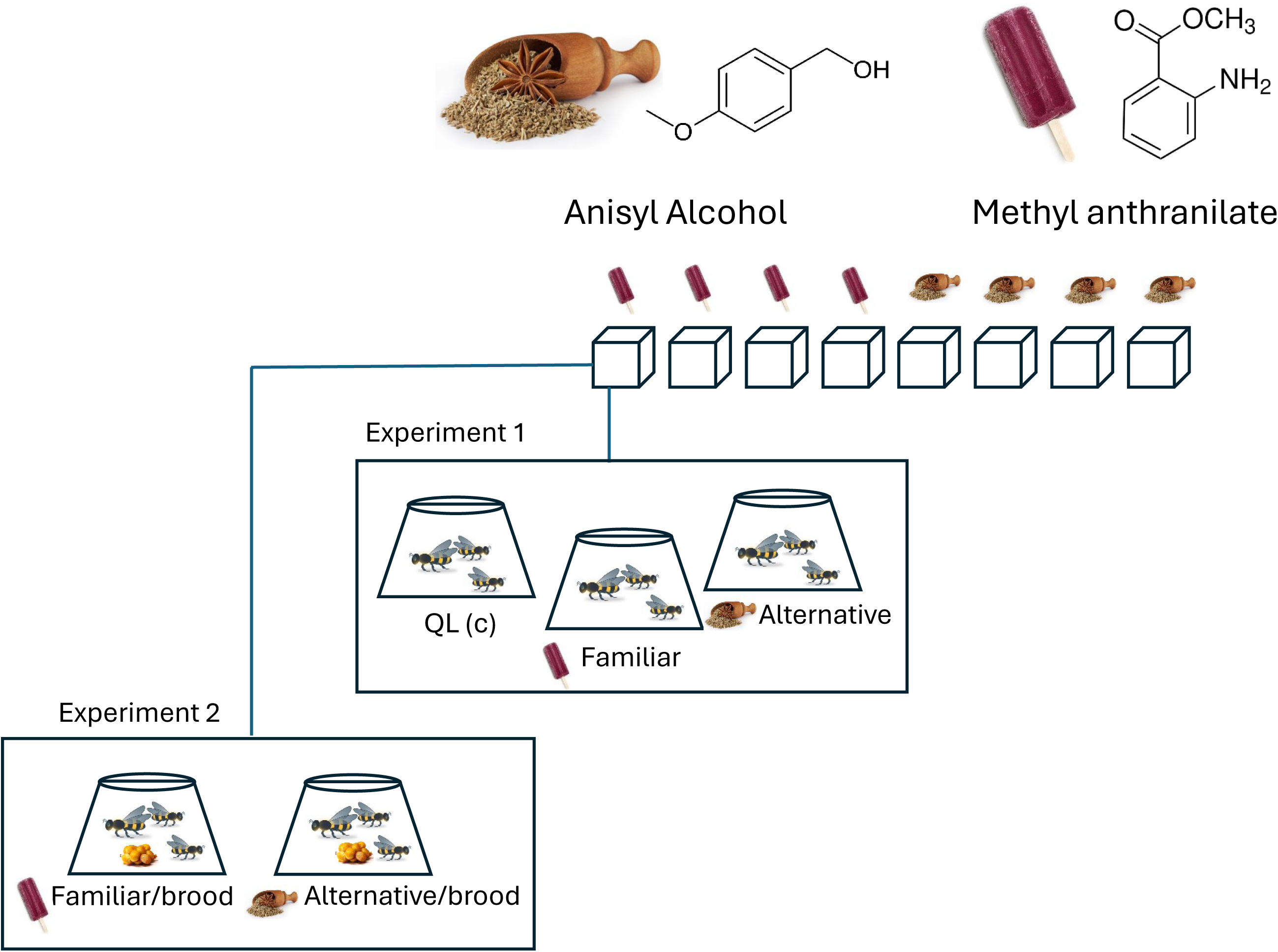
Experimental design. Queens from eight *Bombus terrestris* colonies were treated daily with 1 mg of either anisyl alcohol or methyl anthranilate shortly after the emergence of the first worker. Once colonies reached 20 workers, nestmate bees were sampled progressively in groups of three and exposed to either the maternal colony’s odor (familiar) or an alternative odor. Queenless control groups received a scentless cotton wick. Additional queenright workers were sampled to assess colony reproductive status (not shown in the diagram). In the second experiment, groups of three nestmate workers were exposed to either the familiar or alternative odor together with brood from their maternal colony.

### Experiment 1

Cages of three workers were randomly assigned to one of 3 treatments: 1) QL (queenless): these bees were provided with a clean dental wick daily with no treatment. These bees served as a positive control, showing the ability of workers to fully activate their ovaries within a week under QL conditions where the queen is absent; 2) QL-familiar. These workers were provided daily with a dental wick treated with 1 mg/1 µl of the same odor as in their natal colony. Workers from AA-treated queens were provided with AA whereas workers from MA-treated queens were provided with MA. These workers served to test whether they have learned the odor applied onto the queen and have recognized it in association with the queen or the nest and consequently are reproductively affected by it; 3) QL-alternative. These workers were provided daily with a dental wick treated with a different odor than in their natal colonies (i.e., workers from AA-treated queens were provided with MA and vice versa). These workers served to test whether workers are reproductively affected by an alternative odor which was not associated with their queen. All cages were kept for 7 days, after which they were scanned for the total number of eggs laid in them. In addition, we collected queenright (QR) bees from the mother colonies (i.e., workers from colonies in which a reproductive queen is present) throughout the experiment and froze them immediately (11-15 bees per colony). These bees served as a negative control to assess the level of worker ovary activation in the natal queenright colonies. They were dissected, and their terminal oocytes’ size was measured.

### Behavioral assays

During the first experiment, 15 workers from each colony were sampled for a behavioral test, examining their preference between the odor they experienced during development versus the alternative odor. Bioassays were conducted using a T-maze made of polycarbonate sheets (Video 1). The endpoint of each arm of the maze included a filter containing each of the two odors at the same concentration as applied to the queen. Each arm was connected into a plastic cage. Each worker was given 10 minutes to select the preferred arm, defined as positive if the worker passed the filter and entered the empty cage without the ability to return to the maze. Workers that did not decide within 10 minutes were defined as non-responsive. The odors were alternate between arms, and the maze was wiped clean prior to every bioassay. To prevent resampling of the same bee in consecutive bioassays, each tested worker was marked with a color and returned to her colony at the end of the test. All tests were conducted within three days during the morning hours, about 20 days after the onset of the treatments of the queens, concurrently with the sampling of bees for the first experiment.

### Experiment 2

Following the completion of the first experiment, the above mother colonies received a day or two without sampling to encourage their growth. On days 31-35 all workers from all colonies were placed in cages in groups of 3 nestmates together with a small amount of brood from their mother colony. The approximate amounts and developmental stages of the brood (eggs, young or old larvae, or pupae) were recorded when placed in the cages, and again, more precisely, at the end of the experiment. Cages were randomly assigned to one of two treatments: 1) QL-familiar/Brood: these were provided daily with a dental wick treated with 1 mg/1 µl of the same odor as in their natal colony; 2) QL-alternative/brood: These workers were provided daily with a dental wick treated with an odor different from the one used in their natal colonies. All cages were kept for 7 days, after which they were scanned for the total number of laid eggs.

### Choice and chemical analysis of odors

The odors in this study were chosen based on several parameters. Both odors were novel for the bees, yet abundant in their natural environment, which ensures in all probabilities that workers can perceive them. Anisyl alcohol (AA, CH3OC6H4CH2OH) and methyl anthranilate (MA, C8H9NO2) are floral scents that are liquid at room temperature and occur in plants pollinated by bees, including bumblebees. However, neither compound has been identified in extracts or head space samples from bumblebees at any developmental stage. In addition, the two compounds have different chemical structures (an alcohol and an ester, both containing aromatic rings) but that differ only slightly in their molecular weight (138.16 and 151.65 gram/mol in AA and MA, respectively), and volatility (boiling point 259 vs. 256, flesh point 110 vs. 104 and vapor pressure at 25° C is the same: 0.0+0.5 for both). To determine the daily concentration and frequency of application, we conducted an additional experiment where we applied 1 mg/1 µl of AA and MA on the thorax of queenless workers kept in small groups and measured their quantity on the thoracic surface at timepoints 0, 24h, 48h and 72h post application. Workers (4-6 per timepoint, n=42) were frozen at -80° C. Their thorax was then immediately separated from the rest of their body and washed in hexane for 10 minutes. The thorax was gently removed from the solvent, and the extracts were analyzed using an Agilent 7890A GC equipped with a HP-5ms column (0.25id x 30m x 0.25 µm film thickness) connected to an Agilent 5975C mass spectrometer. The run was performed in splitless mode with temperature program from 60 °C at 10 °C/min up to 340 °C. The resulting chromatograms and spectra were analyzed using MSD ChemStation software (Agilent) and all peaks were identified using the NIST database and by comparing retention times and mass fragmentation with synthetic compound standards. Compounds were quantified using two internal standards: decanol and 2 undecanol, added to each extract at a concentration of 0.5 mg/1 µl. As a control, we also quantified three cuticular hydrocarbons constituents (C23, C25 and C27) (Supplementary Information, Fig. S2). Compound quantification was made in comparison with the two internal standards.

### Assessing worker reproduction

Queenright workers from experiment 1 were frozen immediately after sampling and stored at -80°C. Queenright workers do not lay eggs due to the queen’s presence, but they may vary in ovary activation. Therefore, their reproductive status was assessed based on the size of their terminal oocytes ^47,48^. Queenless workers, in contrast, activate their ovaries rapidly and are expected to have fully developed ovaries within one week ^47,49,50^. Indeed, eggs were found in 96% and 76% of the cages in the first and the second experiment, respectively, and altogether in 90% of the cages. Their reproductive status was therefore assessed by counting the number of eggs laid.

### Ovarian activation

Each worker has eight ovarioles, four in each ovary. The three largest terminal oocytes, with at least one measured from each ovary, were measured to the nearest micrometer. The mean size of these three oocytes was used as an index of ovarian activation ^36^.

### Egg laying

The cumulative number of eggs was counted after seven days. Workers typically lay 2–4 eggs per cell, and the cells are often merged (see Supplementary Information, Fig. S1C). We counted the number of eggs per cage.

To examine the combined effect of odor treatment and brood on worker egg laying, we used two matrices for the brood: (1) the total number of brood at the end of the experiment; and (2) the change in the brood developmental stage between the start and end of the experiment. Brood categories were defined as follows: Egg-Larva means that the brood stage at onset was eggs that hatched as larvae during the 7-day experiment, Larva-Larva are larvae who remained so throughout the experiment, and Larva-Pupa are larvae that pupated by the end of the experiment ^27^. Eggs in *B. terrestris* hatch within 5 days, larva development of workers (no queens were developed in the experiment) takes about 10 days, and the pupa stage takes another 10 days ^51^. Any eggs found in the cage at the end of the 7 days were laid by workers, since any queen eggs originally placed in the cage are expected to hatch within the timeframe of the experiment.

### Statistical analysis

All analyses and visualizations were performed using JMP Pro 18. Quantification of cuticular compounds across time points (Fig. 2) was analyzed using a fixed-effects model followed by a Tukey post hoc test. To examine colony growth (Fig. 3a), we used a Linear Mixed Model (LMM) to compare the number of newly emerged workers over time during the growth and plateau phases, with colony ID included as a random effect. Ovarian activation in queenright workers across colonies and time (Fig. 3b) was analyzed using a linear model with colony and colony age as fixed effects. To assess whether workers preferred the odor they were reared with (Fig. 4), we performed Chi-square goodness-of-fit tests comparing the number of workers choosing each odor (AA or MA) against a null expectation of equal preference (50:50), and also examined the combined data using a Generalized Linear Mixed Model (GLMM) with colony ID as a random factor. Non-responding individuals were excluded from these analyses. To test the effect of treatment on the number of eggs laid in Fig. 5, we used GLMM with a Poison distribution. We used cage treatment and colony odor as fixed effects, and used colony ID, colony age at the time of cage setup, and incubator identity (AA or MA) as random effects. To examine the combined effect of brood and odor on egg-laying counts (Fig. 6), we used a GLMM with brood amount or developmental stage, treatment, and their interaction as fixed effects, and colony ID, colony odor, and colony age as random effects. Statistical significance was accepted at α = 0.05.

**Figure 2.**
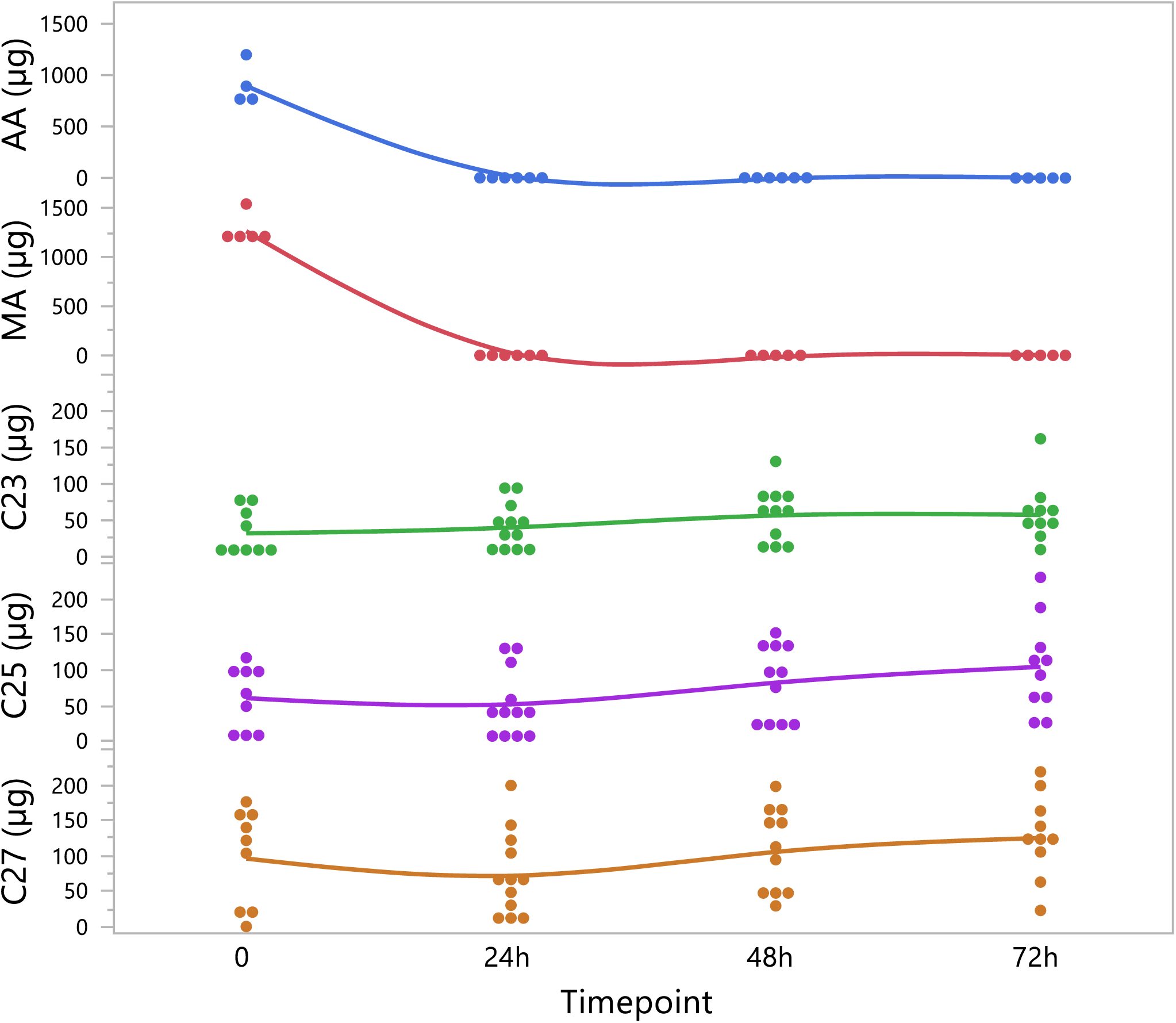
Persistence of experimental odors on the worker cuticle. GC/MS quantification of methyl anthranilate (MA), anisyl alcohol (AA), and three cuticular hydrocarbons (C23, C25, and C27) were quantified over time. Each compound was applied to the worker cuticle at a dose of 1 mg on day 0 and quantified by GC/MS at 0, 1, 2, and 3 days post-application. Quantities were normalized to two internal standards.

**Figure 3.**
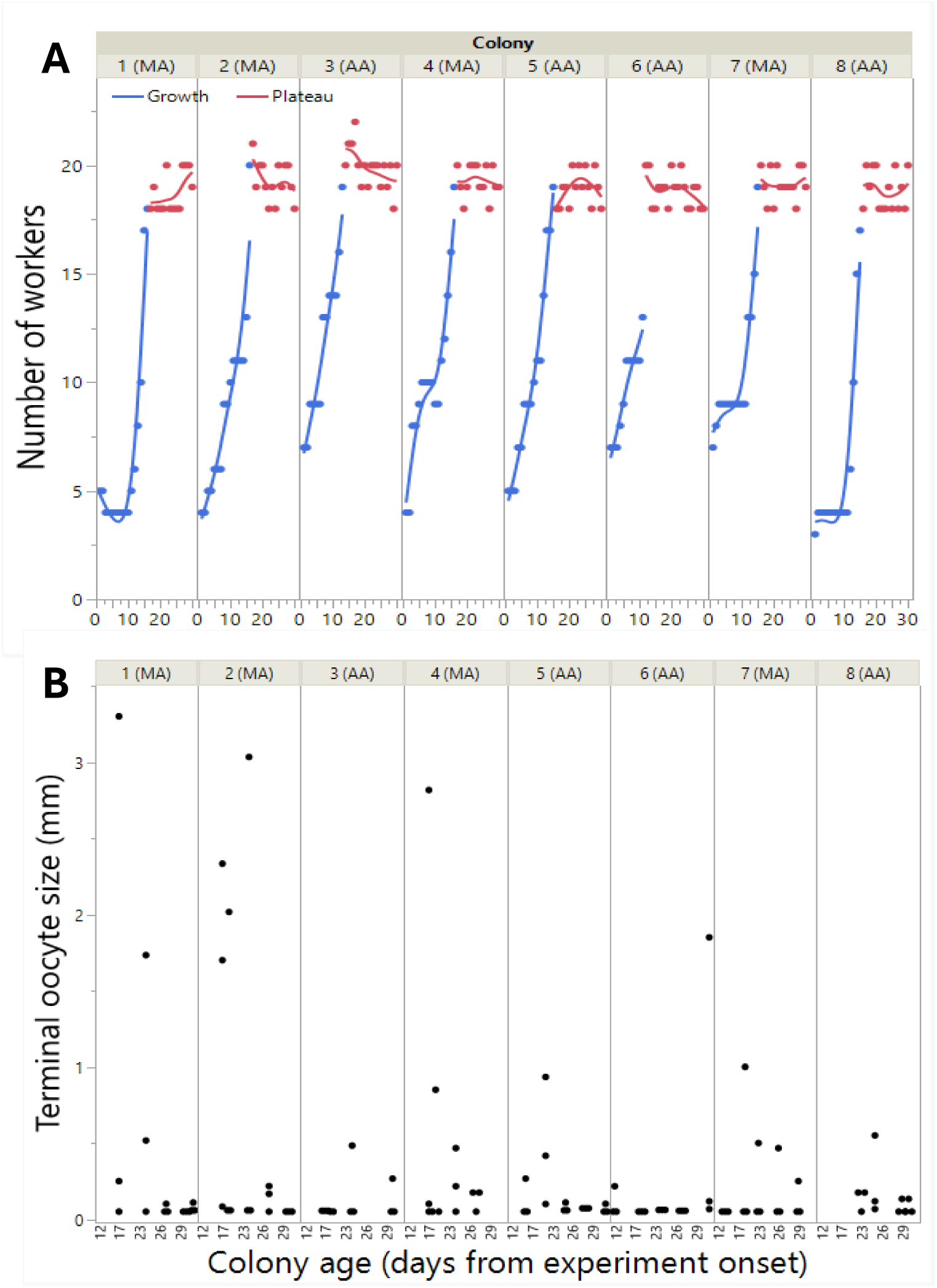
Colony growth and worker ovary activation throughout the experiment. Colony growth and worker ovary activation are shown for the eight colonies used in the study. Colonies reached 20 workers within 12–22 days of the experiment (growth phase) and were then maintained at 20 workers (plateau phase) (A). Workers were sampled during the plateau phase for experiment 1 (days 12–30) and experiment 2 (days 31–35). To assess worker reproductive status, additional workers were sampled directly from the colonies between days 15–30, and their ovaries were measured immediately (B).

**Figure 4.**
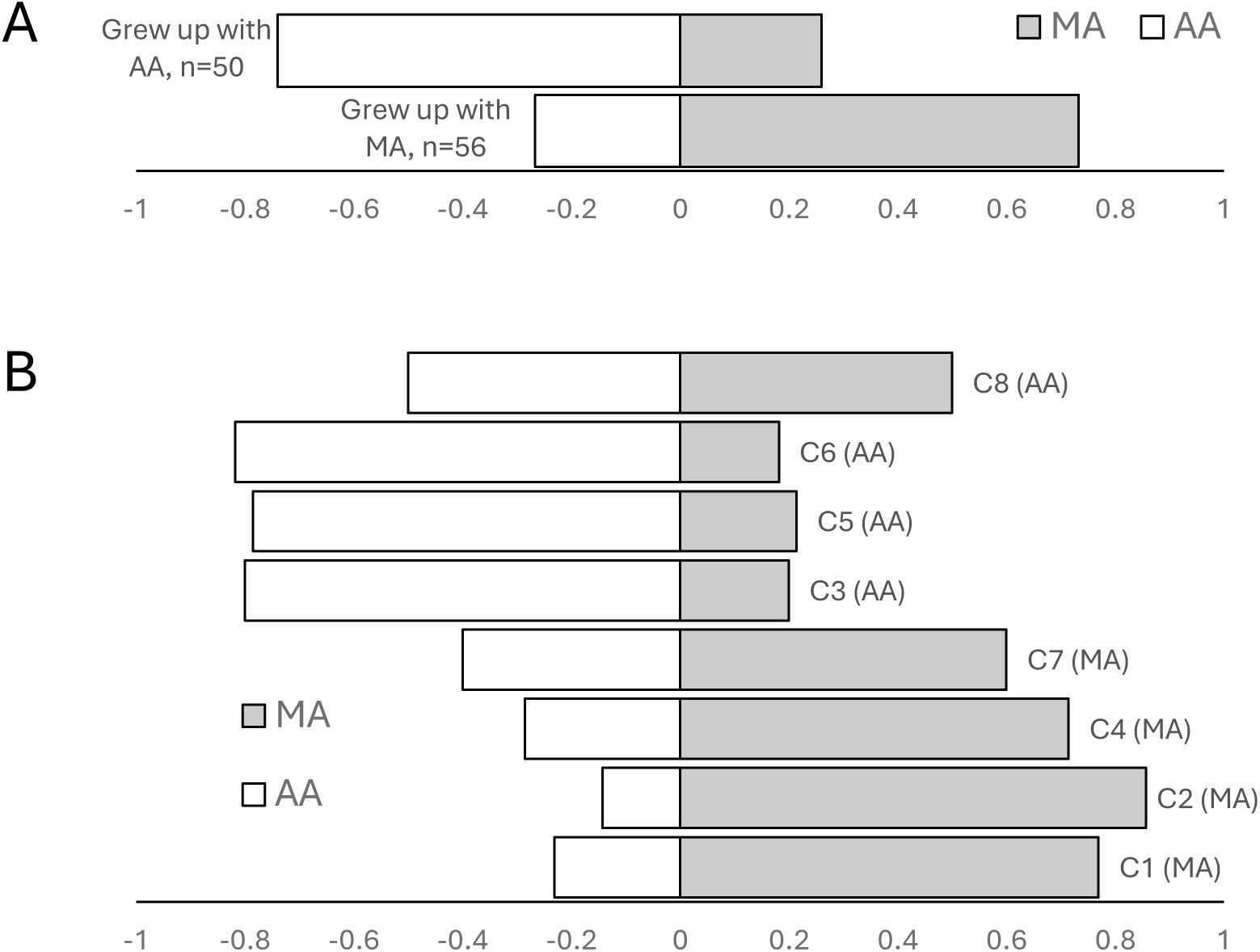
Learned odor preference in worker bees. Two-choice olfactometer bioassays between methyl anthranilate (MA) and anisyl alcohol (AA) showing overall a threefold preference for the queen associated odor (A), with similar patterns across colonies (n = 15 workers per colony, B). Workers were given 10 min to choose before being classified as non-responsive. Non-response rates were low (0–13%) in all colonies except colony 8, where 50% of workers were non-responsive and no clear preference was detected.

**Figure 5.**
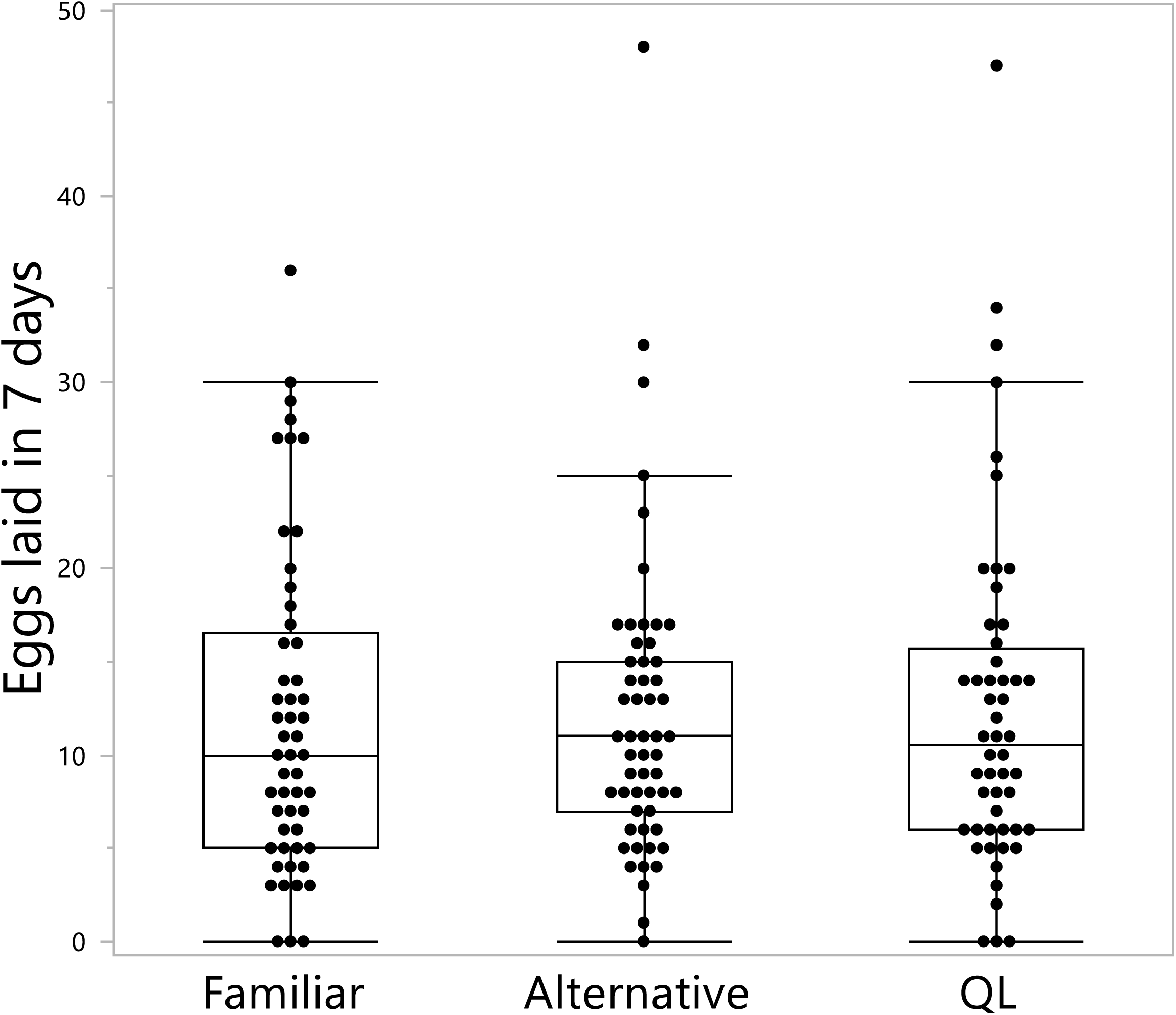
Effects of learned odor on worker egg laying. Worker egg laying following exposure to methyl anthranilate, anisyl alcohol, or no odor. Groups of three nestmate workers were housed together for 7 days and provided daily with a cotton wick treated with 1 mg of either the odor previously associated with their queen (familiar), an alternative odor not previously encountered, or an untreated scentless wick (QL control).

**Figure 6.**
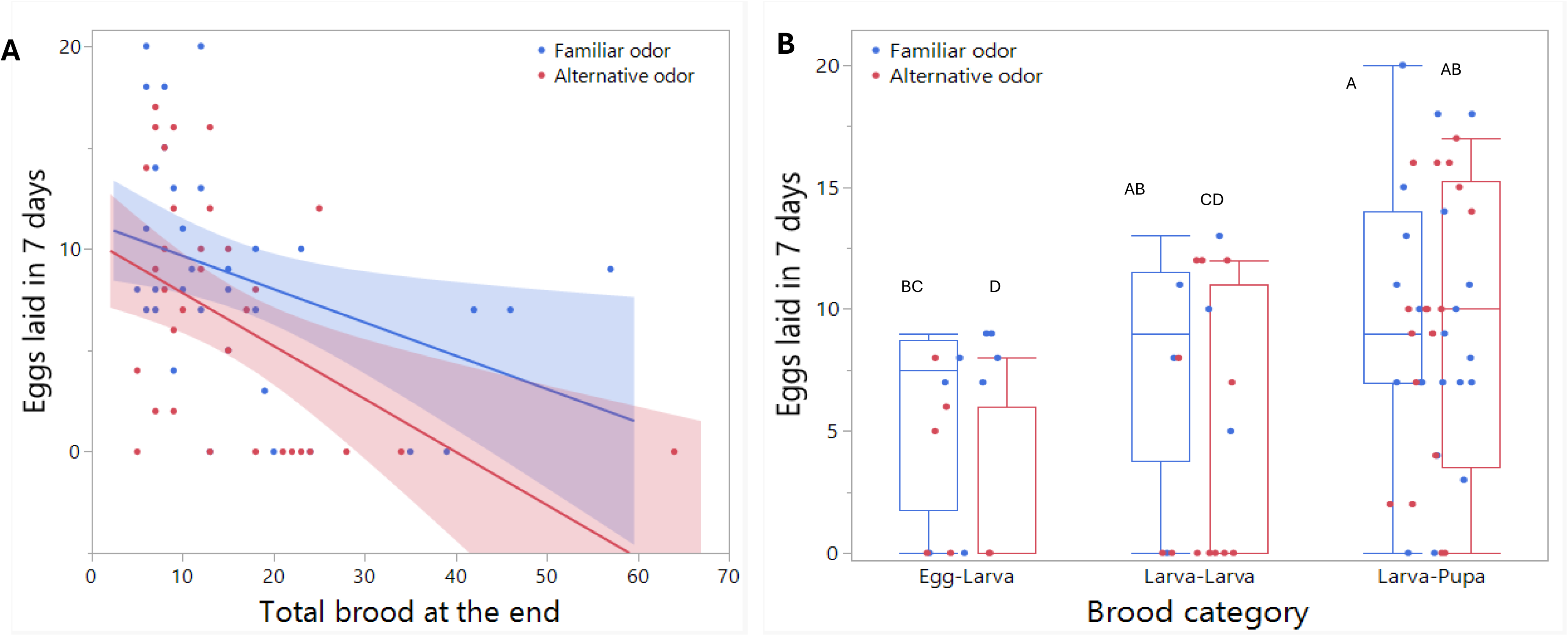
Effects of learned odor on worker egg laying in the presence of brood. Worker egg laying is shown as a function of the odor treatment and the total number of brood present at the end of the experiment (A) and the change in brood developmental stage between the start and end of the experiment (B). Groups of three nestmate workers were housed for 7 days with brood from their natal colonies and provided daily with a cotton wick treated with 1 mg of either the odor previously associated with their queen (familiar) or the alternative odor. Different letters above the columns in (B) indicate significant differences at α = 0.05.

## Results

Quantifying AA, MA and the three major cuticular hydrocarbon constituents on workers’ thoracic cuticle show that the two odors used in the study reached nearly zero amount within 24 hours from application (AA: F_3,17_=120.1, p<0.001; MA: F_3,17_=379.9, p<0.001; both followed by Tukey post-hoc test p<0.001 for timepoint 0 vs. all other timepoints) as opposed to the amounts of the three cuticular hydrocarbons that remain similar over time (C23: F_3,38_=1.24, p=0.3; C25: F_3,38_=2.11, p=0.11; C27: F_3,38_=1.46, p=0.23) (Fig. 2).

Therefore, we chose to apply 1 mg/1 µl onto the thoracic cuticle of queens daily.

All colonies developed normally, reaching 20 workers within 12–22 days (Fig. 3A). There was a significant effect of colony age on worker number during the growth phase (F_1,108.4_=274.9, p < 0.001), indicating a consistent increase in population across colonies. No such effect was observed during the plateau phase (F_1,116.6_=1.82, p=0.17), when colonies were maintained at 20 workers. The ovaries of queenright workers were inactive, and oocyte size did not vary over time (F_14,88_=1.21, p=0.28) or across colonies (F_7,88_=1.16, p=0.32) (Fig. 3B), indicating that all colonies remained during the pre-competition phase during sampling.

Preference bioassays between the two odors showed that workers generally favored the odor they were reared with. When all colonies were pooled together (GLMM with colony ID as a random factor), workers were 3 times more likely to prefer the odor they experienced during development over the alternative odor (74% vs 26%). The effect was highly significant (GLMM, F_1,5.1_=21.9, p=0.005) and the random effect of colony accounted for <1% of the total variance in the response and was insignificant (Wald p-value = 0.96) (Fig. 4A). Among colonies treated with MA, significant preferences for MA were observed in colony 1 (χ² = 3.77, p = 0.05) and colony 2 (χ² = 7.14, p = 0.007). In colony 4 and 7, there was a trend to prefer the queen odor, but the preference was not statistically significant (χ² = 2.57, p = 0.11; χ² = 0.60, p = 0.44, respectively). Among colonies treated with AA, significant preferences for AA were found in colonies 3 (χ² = 5.40, p = 0.02), 5 (χ² = 4.57, p = 0.03), and 6 (χ² = 4.45, p = 0.03), but not in colony 8 (χ² = 0, p = 1.00) (Fig. 4b). A clear behavioral response to the learned odor is further shown in Video 2, which captures an immediate, stereotypical excitation of workers following the introduction of the odor into the colony on a dental wick.

The number of eggs laid within 7 days in the absence of brood did not differ between QL workers that were exposed to the familiar odor, the alternative odor, or were not exposed to any of the two odor treatments (GLMM, F_2,156_=0.84, p=0.43) (Fig. 5). The average number of eggs laid within 7 days in all three groups was 12. The colony odor had no effect either (F_1,6_=0.009, p=0.92). The random factors colony ID and incubator had no impact on the model’s results. The random factor ‘colony age’ was significant (Wald p-value= 0.01) and accounted for 19% of the variation. However, there was no correlation between the colony age and the number of eggs laid by workers (r=0.13. p=0.1, n=160).

In Fig. 6, running GLMM with poison distribution, two fixed effects and 4 random factors to analyze the effect of treatment and brood on egg laying didn’t converge due to complexity. Thus, we tested the effect of the random factors one by one and included them in the model only if they had a significant effect. The effect of treatment and amount of brood with ‘colony’ as a random factor showed that both were significant, but colony had no significant effect on the number of eggs laid (Wald p-value = 0.09) and accounted for only 14% of the variation. Running the same model with ‘colony age’ as a random factor showed a significant effect of treatment and brood amount but not of colony age (Wald = 0.26, 8% of the variation). The same was obtained for the random factor ‘incubator’ (Wald = 0.58, 1.3% of the variation), and the random factor ‘colony treatment’ (Wald = 0.98, <0.01 of the variation). Overall, none of the random factors justified their addition into the model. When the model run without random factors, the number of worker-laid eggs within 7 days in the presence of brood was significantly affected by the treatment (GLMM, χ² = 26.4, p < 0.001), the amount of brood (χ² = 68.3, p < 0.001) and the interaction between the two was significant (χ² = 20.9, p < 0.001). Overall, brood amount negatively affected the number of eggs laid, and workers exposed to a familiar odor laid more eggs compared to the workers exposed to the alternative odor (Fig. 6A). Finally, running the model considering the developmental stage of the brood throughout the experiment instead of the amount of brood resulted in the same outcomes with a significant effect of the treatment (χ² = 17.1, p < 0.001), the brood category (χ² = 47.9, p < 0.001) and the interaction between them (χ² = 10.7, p = 0.004). Overall, young brood (i.e., eggs that became larvae or young larvae) negatively affected the number of eggs with workers laying more eggs when exposed to the familiar odor (Fig. 6B). No differences in the number of eggs laid by workers were found between the odors when the brood was old (i.e., pupae)

## Discussion

Pheromonal responses in insects are generally assumed to be innate and hard-wired, but a growing body of evidence shows that the response to pheromones can be shaped by learning or experience ^9^. In this study, we hypothesized that perfuming the queen with an external odor will presumably create an association between her pheromonal blend and the new odor. Accordingly, we tested whether the learned odor alone could represent the queen’s presence and influence worker behavior and physiology. Workers reared with a perfumed queen preferred the applied odor over the alternative odor in a T-maze, indicating that they can learn a preference for an external odor. However, it remains unclear whether the odor was learned and encoded as a general nest odor after diffusing into the colony environment, or whether it was integrated in the workers’ brains as a component of the queen pheromone. Additionally, when workers were exposed to the learned odor in the context of brood, they increased their overall egg laying. This demonstrates an associative learning with an external odor, but the change in reproductive behavior was opposite to what would be expected from a compound representing the queen ^1,21^. Below, we discuss possible explanations for these findings.

If workers associated the odor with the queen’s presence and interpreted it as a component of fertility or dominance signal, we would expect it to suppress worker egg laying, but the opposite occurred. A comparison with a previous study ^21^ may suggest a possible explanation to this finding. The previous study showed that worker reproduction in *Bombus impatiens* was fully reduced only in the combined presence of queen pheromone, brood, and a newly-emerged gyne (=a visual cue). Specifically, the effect of three combinations of cues and their effect on worker reproduction were tested: (1) queen pheromonal secretion alone, which did not result in inhibited reproduction in workers; (2) queen pheromonal secretion and a newly-emerged gyne, which partially inhibited reproduction, and (3) queen pheromonal secretion, newly-emerged gyne, and brood, which fully suppressed worker reproduction.

The study did not test the effect of queen secretion and brood alone (like in the current study). It is plausible that, had this been tested in the previous study, reproduction would not have been suppressed because of the absence of the visual cue provided by the gyne. The absence of a visual cue in the present study may have signaled to workers that the colony was entering the competition phase ^25^, when the queen may be absent or compromised while queen odors and brood remain present, thereby triggering worker reproduction before the end of the season. Visual stimuli have been shown to be important in bumblebees, and their absence leads to increased volumes of the mushroom body lobes and calyces ^52^. Alternatively, workers may have learned an association between the odor and the nest, and upon separation from the queen, they associate the odor with the sanctity and resource richness of the nest, which could encourage them to initiate reproduction. While the later explanation (odor equals to nest) is based upon contextual learning in bees which has been demonstrated numerous times ^16,17^, the first explanation (odor equals to queen) would involve individual specific learning, which to our knowledge has not been shown before. Further research is needed to test these hypotheses. Such studies could distinguish between these possibilities by applying the same odor to a dummy queen or to nest material alone and testing for similar behavioral responses, or by applying different odors to the queen and nest and assessing which workers learn and respond to. Another valuable control would be to include untreated colonies and test their preference and responses to the treatment odors. Applying odors at different doses or with different chemical structures (i.e., less volatile), as well as testing worker responses to the queen cuticular lipids, would further clarify whether volatility plays an important role in this learning and whether it masks the original odor of the queen.

Another factor that may influence worker learning capacity is age, as learning tends to be more efficient in younger bees ^52,53^, but associative learning may become stronger when exposure is extended. Honey bee workers, for instance, respond to the queen mandibular pheromone only when young, while older workers tend to ignore it ^54,55^. In the present study, workers were of randomly mixed ages, which may have resulted in learning variation and consequently variation in reproductive decision. We tend to exclude this possibility since colony age did not affect the number of eggs workers laid. There was no correlation between colony age and egg production, as would be expected if older colonies contained older workers more ready to reproduce. Eggs were laid in most cages (90%), and the proportion of cages with eggs was actually higher in the first early experiment (96%) than in the second (75%). This difference may be explained by brood presence and developmental stage, with younger and more abundant brood exerting stronger inhibition on egg laying. Overall, workers were attracted to the odor applied to their queen and laid more eggs in its presence, and these behaviors were independent of any random effects recorded in the study, excluding the brood, but the underlying mechanism for this attraction remained unresolved.

The learned odor likely has a releaser function, inducing an immediate behavioral change ^56^. The immediate, stereotypical excitation of workers in response to the learned odor is further demonstrated in Video 2. The odor altered worker attraction and egg-laying behavior but was unlikely to affect ovarian activation (i.e., the size of oocyte size), as most workers had fully activated ovaries and eggs were laid in most cages regardless of treatment. However, treatments differed by the number of eggs laid in them. Although egg laying requires fully activated ovaries, it involves muscle contraction and is regulated separately from oogenesis and thus may also be considered a releaser effect.

Finally, it has been suggested that pheromones promoting reproductive behavior carry intrinsic reward value, which can be overridden by experience - particularly through associative learning. While many such modification in pheromonal responses have been documented in mammals ^9^, insects have received far less attention, largely due to the assumption that their responses to pheromones are instinct-driven^16^. However, social insects offer an excellent model to explore the flexibility of pheromonal responses. Their social organization creates strong asymmetries in fitness outcomes - where most individuals forego reproduction gaining only indirect fitness benefit, while a few gain direct and more substantial fitness benefits. This imbalance may favor the evolution of learning mechanisms that modify pheromonal responses and thus provides an ideal system to test the role of experience and learning in pheromone-guided behavior.

## Acknowledgment

We would like to thank Dr. Eran Levin for providing the space and the incubators and for Dr. Inon Scharf for providing the T-maze. We also thank the four anonymous reviewers whose comments greatly improved the manuscript.

## Data Availability

The source data supporting the findings of this study are provided in Supplementary Data 1.

## Author Contributions

EA and AH designed the study and wrote the manuscript. EA conducted the experiments and performed the analyses. AH conducted the chemical analysis.

