## Supplementary information for "A social bee (*Bombus terrestris*) learns to associate a novel odor with social context"

**Table S1.** The overall sample size used in the study split by experiment, treatment and colony and the total number of workers produced by the colonies throughout the experiment.

| Treatment | Unit of replication | Col. 1 | Col. 2 | Col. 3 | Col. 4 | Col. 5 | Col. 6 | Col. 7 | Col. 8 | Total cages | Total workers |
|---|---|---|---|---|---|---|---|---|---|---|---|
| Colony odor | | MA | MA | AA | MA | AA | AA | MA | AA | | |
| QR workers | Workers | 15 | 15 | 11 | 12 | 15 | 15 | 15 | 12 | | 110 |
| Bioassays | Workers | 15 | 15 | 15 | 15 | 15 | 15 | 15 | 15 | | 120 |
| Experiment 1/QL | Cages | 5 | 7 | 7 | 7 | 6 | 6 | 8 | 6 | 52 | 156 |
| Experiment 1/Familiar | Cages | 6 | 7 | 7 | 5 | 7 | 6 | 9 | 6 | 53 | 159 |
| Experiment 1/Novel | Cages | 7 | 8 | 6 | 7 | 7 | 6 | 9 | 5 | 55 | 165 |
| Experiment 2/Familiar_brood | Cages | 6 | 6 | 5 | 4 | 5 | 2 | 6 | 3 | 37 | 111 |
| Experiment 2/Novel_Brood | Cages | 6 | 6 | 5 | 4 | 4 | 2 | 7 | 3 | 37 | 111 |
| Total workers produced by colony | | 120 | 132 | 116 | 108 | 117 | 96 | 147 | 96 | | |

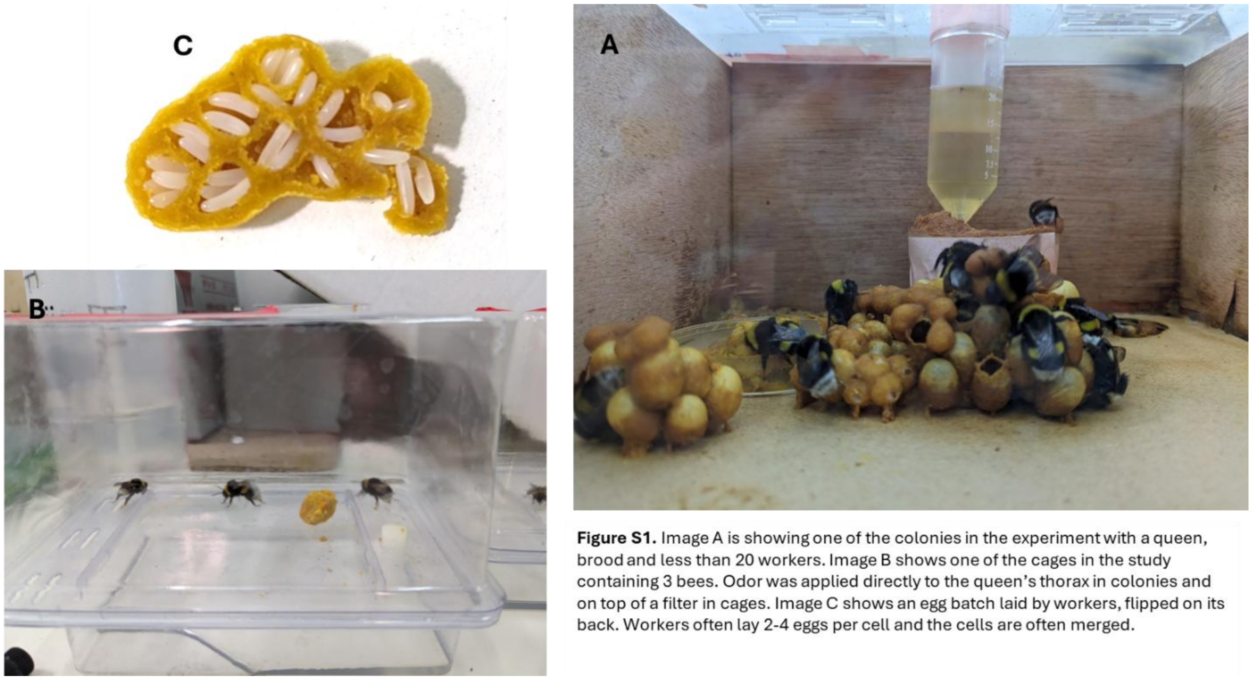


**Figure S1.** Image A is showing one of the colonies in the experiment with a queen, brood and less than 20 workers. Image B shows one of the cages in the study containing 3 bees. Odor was applied directly to the queen's thorax in colonies and on top of a filter in cages. Image C shows an egg batch laid by workers, flipped on its back. Workers often lay 2-4 eggs per cell and the cells are often merged.

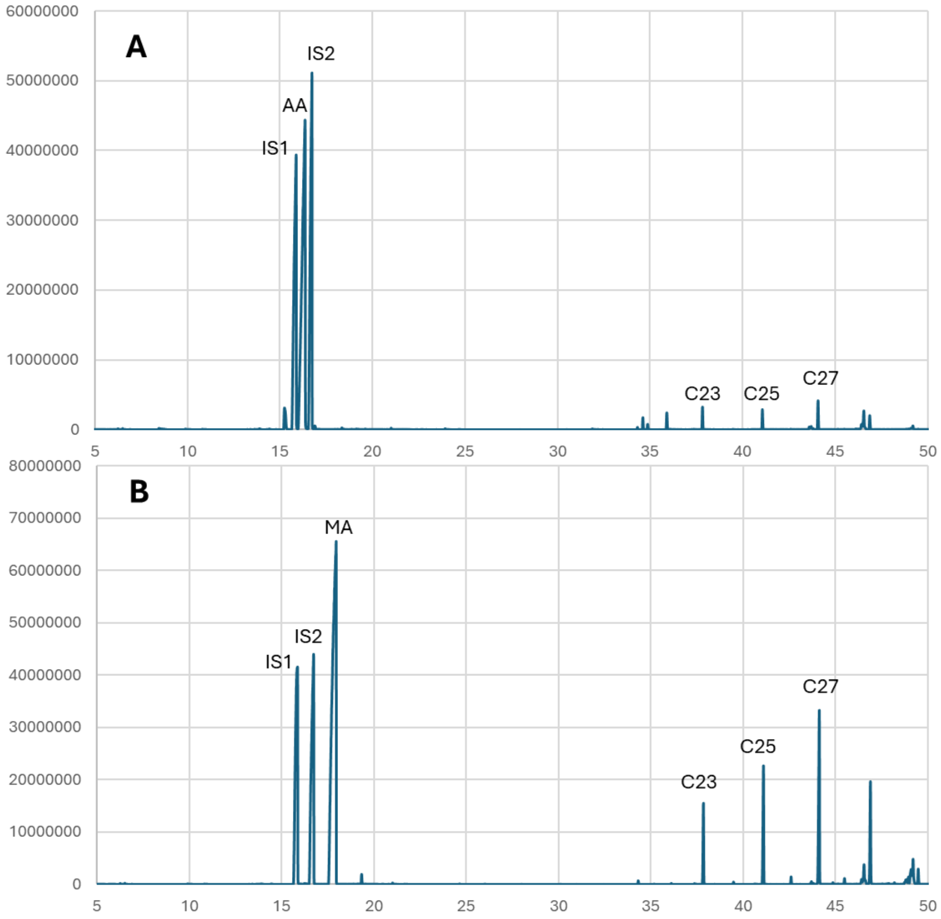


**Figure S2.** GC/MS chromatograms of 1 mg anisyl alcohol (A) and methyl anthranilate (B) applied to the cuticle of a worker bee. Three common hydrocarbons (C23, C25, C27) were also quantified. Decanol (IS1) and 2-undecanol (IS2), were used as internal standards for compound quantification (0.5 mg per sample, each).
